# Effect of Initial pH and Organic Matter Concentration on Production of Volatile Organic Acids Through Acidogenesis of Sugarcane Vinasse

**DOI:** 10.1101/2021.02.22.432232

**Authors:** Juliana Martins, Hugo Valença de Araújo, Gustavo Mockaitis, Ariovaldo José da Silva

## Abstract

Sugarcane vinasse is an industrial liquid waste generated in great amounts in Brazilian ethanol industries. Nowadays its main use occurs at sugarcane crops, where vinasse is applied as a nutrient source for fertirrigation. However, continued use of vinasse in soil can cause several environmental impacts. So, aiming to provide a more environmentally friendly destination to the effluent, the goal of this work was to investigate the acidogenesis using a synthetic vinasse as substrate, focusing on the effects of initial pH and Chemical Oxygen Demand (COD) on short chain organic acids (SCOAs) concentrations. Synthetic vinasse was prepared at laboratory taking some real sugarcane vinasse composition given in previous works as references. So, major contribution presented here is the investigation on obtaining high added-value SCOAs from a simulated effluent. Cattle manure sludge was utilized as inoculum to promote the conversion of carbohydrate (sucrose, Suc) in synthetic vinasse into SCOAs in batch reactors during a total incubation time of 72 h. Acidogenesis profiles have shown that concentration of lactic acid (HLa) was prevailing among all metabolites, indicating that process followed through an essentially lactic route. Furthermore, considerable concentrations of propionic, acetic and isobutyric acids were also verified at some specific operation times, while solventogenesis was not detected at all. The greatest peak of lactate content was 4.96 g HLa L^−1^, observed under initial pH 6.0 and 25 g COD L^−1^, at 16 h. Maximum of lactate productivity was 332.10 g HLa L^−1^ h^−1^ at 8 h, associated to a yield of 189.14 g HLa (g Suc)^−1^, under initial pH 7.5 and 20 g COD L^−1^.

## 1. INTRODUCTION

Generation of agroindustrial wastes in great amounts has drawn attention to the final disposition given to them, which should be realized in such a way to minimize impacts on receiving bodies since effluents generally present high pollution potential. In sugar and ethanol production process from sugarcane, both products of high economic importance in countries such as Brazil, two main residues are obtained, namely sugarcane bagasse and vinasse. Thus, renewability of ethanol fuel depends strongly on the destination these waste materials receive. Since a long time sugarcane bagasse has been successfully utilized in generation of steam and electricity by means of its burning in boilers existing in sugar and alcohol industrial units (DIAS *et al*., 2015). Vinasse, although traditionally applied as a nutrient source for fertirrigation in sugarcane crops, can cause severe environmental problems mainly if its use is continued, so that unconventional processing of it have been the target of several works more or less recently (HOARAU *et al*., 2018)

It is expected that ethanol production will grow in the coming years and reaches 134 billion of liters in 2024, with United States, Brazil, European Union and China as the world’s largest producers (OECD, 2015). In 2018, United States was responsible for 56% of all ethanol generated in planet. Also in 2018, production was equivalent to about half of the American in Brazil (28%) (RFA, 2019), highlighting the importance of Brazilian ethanol in the global context. So, vinasse is indeed a relevant agroindustrial waste in Brazil, generated as a liquid by-product in hydrous ethanol purification system. It is an acidic (pH 3.5-5.0), dark brown coloured and unpleasant odour stream with a high organic load, implying a high Chemical Oxygen Demand (COD) (HOARAU *et al*., 2018).

Some previous studies has already pointed out that use of vinasse as a fertilizer may have positive consequences for sugarcane crops (BRITO *et al*., 2009; JIANG *et al*., 2012; UYEDA *et al*., 2013; PREVINA & SARAVANAM, 2013; FUESS e GARCIA, 2014), which is its main use nowadays. However, Fuess and Garcia (2014) highlighted some important disadvantages, mostly observed if vinasse is continuously disposed on soil, namely salinization and sodification, organic overloading, permanent acidification of water bodies, interference in the photosynthesis of aquatic plants and inhibition of seed germination. These negative effects has motivated strongly the search for alternative processes aiming the degradation of sugarcane vinasse.

Sugar and ethanol factories generate an average of 156 L of vinasse and 250 kg of bagasse for each ton of sugarcane processed, providing 12 L of ethanol and 94 kg of sugar (sucrose). Therefore, an average ratio of 13 L of vinasse by liter of ethanol is attained (GUNKEL *et al*., 2006), which can varies widely as a function of production process parameters (feedstock quality, ethanol fermentation yield, and so on). Based on these data, perspective is that 1,742 billion of liters of vinasse are to be generated until 2024 (HOARAU *et al*., 2018). Such a great volume of agroindustrial waste is very significant and various methods to convert it into value-added products, especially chemical compounds and energy carriers, have been investigated (FUESS *et al*., 2020; MORAES et al., 2019).

In the context of SCOAs biological production using vinasses resulting from ethanol generation from sugarcane, sugar-beet and other agricultural feedstocks (or synthetic vinasses in order to simulate the real ones), lactic acid (also named lactate) is certainly the most focused metabolite because of its great versatility, since it can be used as an acidifying and flavouring agent, leather softener, antimicrobial preservative and chemical input. A highlighting application of lactic acid occurs as a precursor for the synthesis of polylactic acid polymers in pharmaceutical and plastic industries due to its favourable attributes of biodegradability, biocompatibility and elasticity (HOARAU *et al*., 2018).

Many microorganisms species and processes have been applied to produce lactic acid from vinasse as substrate. Liu *et al*. (2012) used a method based on combined application of microwave and NaOH pre-treatments in order to reduce contents of hemicellulose and lignin in vinasse, resulting in a higher lactate yield in comparison to the case in which each pre-treatment was executed separately. Lactic fermentation of vinasse by zeolite-immobilized cells of *Lactobacillus rhamnosus* ATCC 7469 was performed by Djukić-Vuković *et al*. (2013), which concluded that support material used was effective to promote the process. Similarly, Djukić-Vuković *et al*. (2016) employed the same microbial strain, but using a Mg-modified zeolite to cell retention, also obtaining considerable high yields and productivities in batch experiments.

Djukić-Vuković *et al*. (2015) analysed lactate production from vinasse by batch fermentation with *Lactobacillus rhamnosus* ATCC 7469, reaching higher reaction performance parameters in comparison to other residual substrates, even though vinasse has not been supplemented with nitrogen and other materials. Additionally, as process by-product, Djukić-Vuković *et al*. (2015) obtained a dry solid suitable for feeding of monogastric animals. Employing microorganisms of a mixed culture from an Upflow Anaerobic Sludge Blanket (UASB) reactor and immobilized over particles of a Greek volcanic rock and γ-alumina, Lappa *et al*. (2015a) and Lappa *et al*. (2015b) detected lactate as the prevailing metabolite in continuous acidogenesis tests with vinasse as feed stream.

Moraes *et al*. (2019) claimed the existence of a lack in technical literature concerning works exploring the production of SCOAs and alcohols by vinasse anaerobic digestion. In addition to lactic acid, acetic and butyric acids (acetate and butyrate) have also been synthesized via acidogenesis using vinasse as substrate (LAPPA *et al*., 2015a; MORAES et al., 2019; FUESS *et al*., 2020). Butyric acid finds applications as precursor for biological fermentative production of butanol by *Clostridium spp*. and also as raw material for food, beverages, chemical, textile and pharmaceutical industries (ZHANG *et al*., 2009). However, number of works dealing with SCOAs and alcohols production by vinasse anaerobic digestion is indeed scarce to the present date probably due to the strong emphasis that has been given almost exclusively to hydrogen and methane (H_2_ and CH_4_) generation, which can be executed by means of more mature and practical biotechnological processes (BAGHCHEHSARAEE *et al*., 2008; MORAES *et al*., 2015; FUESS e ZAIAT, 2018).

In this work, the main goal was to carry out acidogenesis batch experiments to evaluate effects of initial pH and COD of a synthetic vinasse on the production of SCOAs. A synthetic vinasse prepared in laboratory was used because it allows a better control of substrate parameters aiming to verify more easily the operating conditions leading to maximization of desirable metabolites contents. By doing this, great variability of real vinasse characteristics, which is typical for the industrial waste, was avoided. Thus, major contribution presented here is the investigation on obtaining high added-value SCOAs from a simulated effluent that has been processed mostly focusing on the production of biogas for use as biofuel.

## 2. MATERIAL AND METHODS

### 2.1 Experimental design

Volatile acids production experiments were performed in 500 mL Duran© flasks, in which 300 mL was working volume, while 200 mL was taken as headspace. Reactor bulk was constituted by 270 mL of the corresponding media and 30 mL of inoculum at 24.33 g TVS (Total Volatile Solids) L^−1^. Therefore, initial inoculum concentration was kept constant in all experiments at 2.4 g TVS L^−1^.

Flasks were maintained in an orbital shaker-incubator to be subjected to a constant rotational speed of 100 min^−1^, under controlled temperature (37 °C). Initial pH of each medium was adjusted by adding sodium bicarbonate (NaHCO_3_). To ensure an anaerobic condition to all experiments, nitrogen was fluxed at constant flow rate in all flasks for 10 min. All experiments were conducted for 72 h and samples of media were collected for monitoring of reactors every 8 h. Table 1 shows the experimental design used in this study.

**Table 1 –.** Matrix of experiments for evaluating the effects of initial pH and COD on SCOAs and alcohols production.

| Reactors | COD<br>(g O<sub>2</sub> L<sup>-1</sup>) | pH |
| --- | --- | --- |
| B1 | 20 | 4.5 |
| B2 | 30 | 4.5 |
| B3 | 20 | 7.5 |
| B4 | 30 | 7.5 |
| B5 | 25 | 6.0 |
| B6 | 25 | 6.0 |
| B7 | 25 | 6.0 |

### 2.2 Media

Media was prepared considering the organic matter composition as defined by Godoi *et al*. (2017) and the characterization of the vinasse from sugarcane as determined by Moraes *et al*. (2019). A complex synthetic media was used in this study rather than real vinasse due to the variation of composition and concentration in real vinasse could undermine the conclusions. However, the media used in this work shows similarity with real vinasse. Table 1 shows the concentration and composition of each media used in this experiment.

**Table 1 –.** Compositions of synthetic media for different COD_sol_.

| Component | Concentration |  |  |
| --- | --- | --- | --- |
| $COD_{sol}$ (g $O_2$ L<sup>-1</sup>) | 20 | 25 | 30 |
| Sucrose (g Suc L<sup>-1</sup>) | 12 | 15 | 18 |
| Propionic acid (g HPr L<sup>-1</sup>) | 0.50 | 0.63 | 0.75 |
| Butyric acid (g HBu L<sup>-1</sup>) | 0.70 | 0.88 | 1.05 |
| Acetic acid (g HAc L<sup>-1</sup>) | 1.30 | 1.63 | 1.95 |
| Ethanol (g EtOH L<sup>-1</sup>) | 1.50 | 1.88 | 2.25 |
| Nitrogen (g N L<sup>-1</sup>) | 0.80 | 1.00 | 1.20 |
| Sulphur (g S-SO<sub>4</sub><sup>2-</sup> L<sup>-1</sup>) | 1.00 | 1.25 | 1.50 |
| Potassium (g K L<sup>-1</sup>) | 2.20 | 2.75 | 3.30 |
| Calcium (g Ca L<sup>-1</sup>) | 0.44 | 0.55 | 0.66 |
| Magnesium (g Mn L<sup>-1</sup>) | 0.25 | 0.31 | 0.38 |
| Sodium (g Na L<sup>-1</sup>) | 0.52 | 0.65 | 0.78 |
| Phosphorus (mg P-PO<sub>4</sub><sup>3-</sup> L<sup>-1</sup>) | 70.0 | 87.5 | 105 |
| Phenol (mg L<sup>-1</sup>) | 6.0 | 7.5 | 9.0 |
| Iron (mg Fe L<sup>-1</sup>) | 10.0 | 12.5 | 15.0 |
| Manganese (mg Mn L<sup>-1</sup>) | 3.00 | 3.75 | 4.50 |
| Zinc (mg Zn L<sup>-1</sup>) | 0.40 | 0.50 | 0.60 |
| Copper (mg Cu L<sup>-1</sup>) | 0.30 | 0.38 | 0.45 |
| Nickel (mg Ni L<sup>-1</sup>) | 0.20 | 0.25 | 0.30 |
| Lead (mg Pb L<sup>-1</sup>) | 0.20 | 0.25 | 0.30 |
| Cadmium (mg Cd L<sup>-1</sup>) | 0.08 | 0.10 | 0.12 |

### 2.3 Inoculum

Inoculum used in batch experiments of this work came from an anaerobic reactor treating cattle manure. After collecting, the anaerobic sludge was subjected to an thermal-acidic pre-treatment, based on the procedures and results found by Mockaitis *et al*. (2020). This pre-treatment was performed only select only microorganisms which are capable of performing hydrolysis and acidogenesis, inhibiting methanogenesis.

Sludge thermal-acidic pre-treatment was performed prior inoculation. The water from the sludge was drained using a 1.0 mm mesh sieve and then heated in a water bath at 80 °C for 30 min. Afterwards, sludge was cooled down to environmental temperature (25-30 °C) using an ice bath. After cooling, pH value was set to 3.0 by the addition of hydrochloric acid (HCl) 1.0 mol L^−1^. Subsequently, the inoculum was stored in a refrigeration chamber at 4 °C for 24 h. Finally, sludge pH was increased to 6.98 by the addition of sodium hydroxide (NaOH) 1.0 mol L^−1^.

### 2.4 Analytical methods

The following parameters were analysed for all samples collected from batch reactors (respective analytical methods are in parentheses): pH (4,500-H+ electrometric method) (APHA, 2012), total sugar content (carbohydrate to sucrose – colorimetric method at wavelength λ = 490 nm) (DUBOIS *et al*., 1956) and cellular growth (optical density, OD, directly linked to medium turbidity, considering a wavelength λ = 600 nm). OD analyses were conducted by reading absorbances of samples diluted with distilled water in the proportion 1:5 (BEGOT *et al*., 1996).

SCOAs and alcohols in samples were evaluated by means of High Performance Liquid Chromatography (HPLC, Shimadzu Scientific Instruments©). HPLC was equipped with a degasifier (DGU-20A^3R^), two parallel pumps (LC-20AD), automatic sampler (SIL-20A_HT_), column oven (CTO-20^a^), ultraviolet detector (UV-DAD) executing the reading at λ = 210 nm (SDP-20^a^), refraction index detector (RID-10a) and controller (CBM-20a). Runnings were carried out in an Aminex column HPX-87H (300 mm x 7.8 mm, BioRad©) keeping a fixed oven temperature of 43 °C. Mobile phase was an aqueous solution of sulfuric acid (H_2_SO_4_) 0.005 mol L^−1^ at a flow rate of 0.5 mL min^−1^. Injected volume of each sample (pre-filtered and under pH ~2.0) in HPLC was 100 μL. Identification of peaks and initial analyses of chromatograms were realized by means of LC Solution Shimadzu© software.

### 2.5 Performance analysis

Performance of batch experiments was evaluated based on parameters such as productivity (*P*, mg SCOA L^−1^ h^−1^) and yield (*Y*, mg SCOA (g Suc)^−1^), calculated according to definitions of Eqs. (1) and (2), respectively, where *C_met,t_* is the metabolite concentration at incubation time t (g L^−1^); *C*_*met,t*=0_ is the initial concentration of metabolite (at time *t* = 0); *C_i_* is the initial concentration of carboydrate (g Suc L^−1^); and *t* is the incubation time (h).

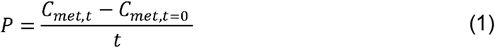

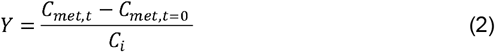

Mass balances in terms of theoretical COD were computed in order to check the consistency of experimental and theoretical results at the end of all batch tests, after 72 h of incubation time. Atomic mass balance equations were developed for each chemical element considering the oxidation reaction by oxygen (O_2_) of 1.0 mol of an organic compound with general molecular formula C_x_H_y_O_z_ representing SCOAs, alcohols and microbial biomass.

The theoretical COD associated to each metabolite (*COD_t,met_*) corresponds to the theoretical quantity of O_2_ required to oxidize completely 1.0 mol of an organic molecule (C_x_H_y_O_z_). Therefore, in order to convert organic compound concentration measured experimentally by HPLC into *COD_t,met_*, it was used the ratio of *m*_O_2__ by *m*_C_x_H_y_O_z__, which are oxygen and organic compound stoichiometric masses, in accordance to Eq. (3). Note that *x, y* and *z* are the numbers of carbon, hydrogen and oxygen atoms existing in organic molecule.

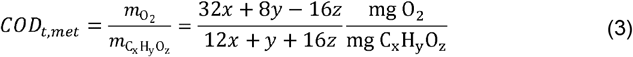

Relations derived from stoichiometric for complete oxidation reactions of organic compounds were employed to compute mass balances referring to detected metabolites [Eq. (3)], sucrose (Suc) and microbial biomass (Bio). Total theoretical COD (*COD_t,T_*) was determined by means of Eq. (4) as a weigthed sum of theoretical CODs (*COD_t,Suc_* for sucrose, *COD_t,Bio_* for biomass and *COD_t,met_* for each metabolite) by correspondent concentrations of all process participants (*C_Suc_, C_Bio_* and *C_met_*, 10^−3^ g L^−1^).

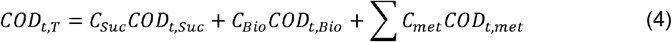

Aiming to verify the compatibility between experimental COD (*COD_exp_*) and total theoretical COD (*COD_t,T_*), a parameter called agreement (*Ag*, %) was calculated as defined by Eq. (5). According to it, the higher the value of *Ag*, the higher the compatibility between both experimental and theoretical CODs.

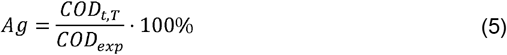

## 3. RESULTS AND DISCUSSION

### 3.1 Acidogenesis profiles

#### 3.1.1 Total carbohydrate concentration, pH, optical density (OD), total SCOAs and lactic acid (lactate) concentrations

Figure 1 shows the time profiles of total carbohydrates concentration (a), pH (b) and OD (c) in different reactional media (B1-B7) through 72 h of processing time.

**Fig. 1 –.**
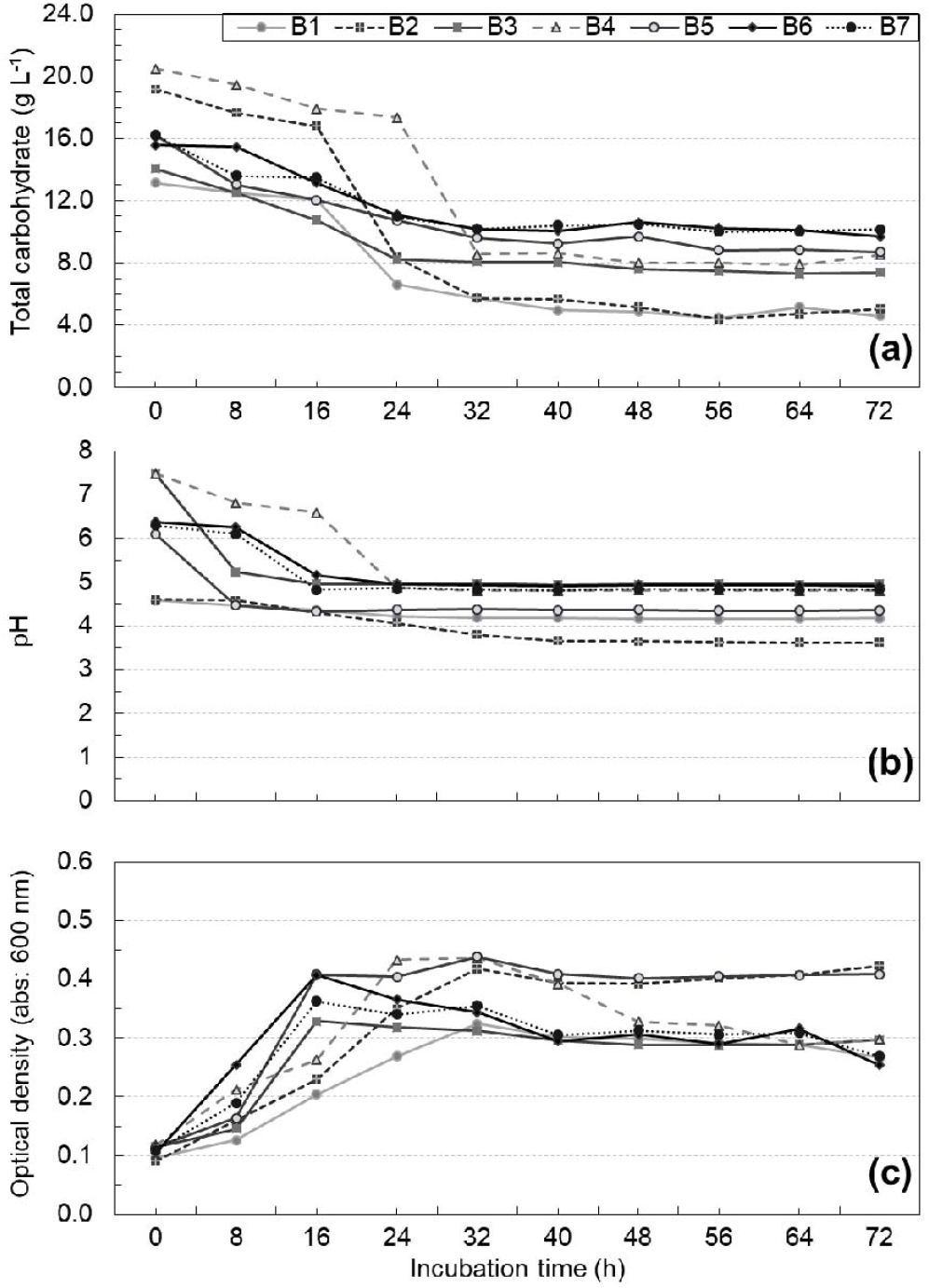
Profiles of monitoring variables in batch reactors during 72 h of total operation: (a) total carbohydrate concentration, (b) pH and (c) optical density (OD).

Carbohydrate consumption occurred mostly before 32 h in all studied conditions, coinciding with the increase in OD profile. This behavior shows that the carbohydrate consumption was associated with cell growth. After 24 hours, the pH reached a stable value for all assays.

Figure 2 shows concentration profiles of total SCOAs and lactic acid (HLac). Note that maximum of concentrations for both total SCOAs and lactic acid are coincident in Figs. 2a and 2b, which is certainly due to lactic acid be the major metabolite. Particularly in Fig. 2b, it is observed that lactate concentration presented increasing profiles between 0 h and 16 h in reactor B5, B6 and B7, attaining maximum of 4.92, 4.66 and 4.96 g HLa L^−1^ at pH 4.3, 5.2 and 4.8, respectively. Probably because of reactors B5, B6 and B7 had been prepared with synthetic vinasse at the same initial pH and COD, trends concerning lactate contents were similar in these media, suggesting a good experimental repeatability of results.

**Fig. 2 –.**
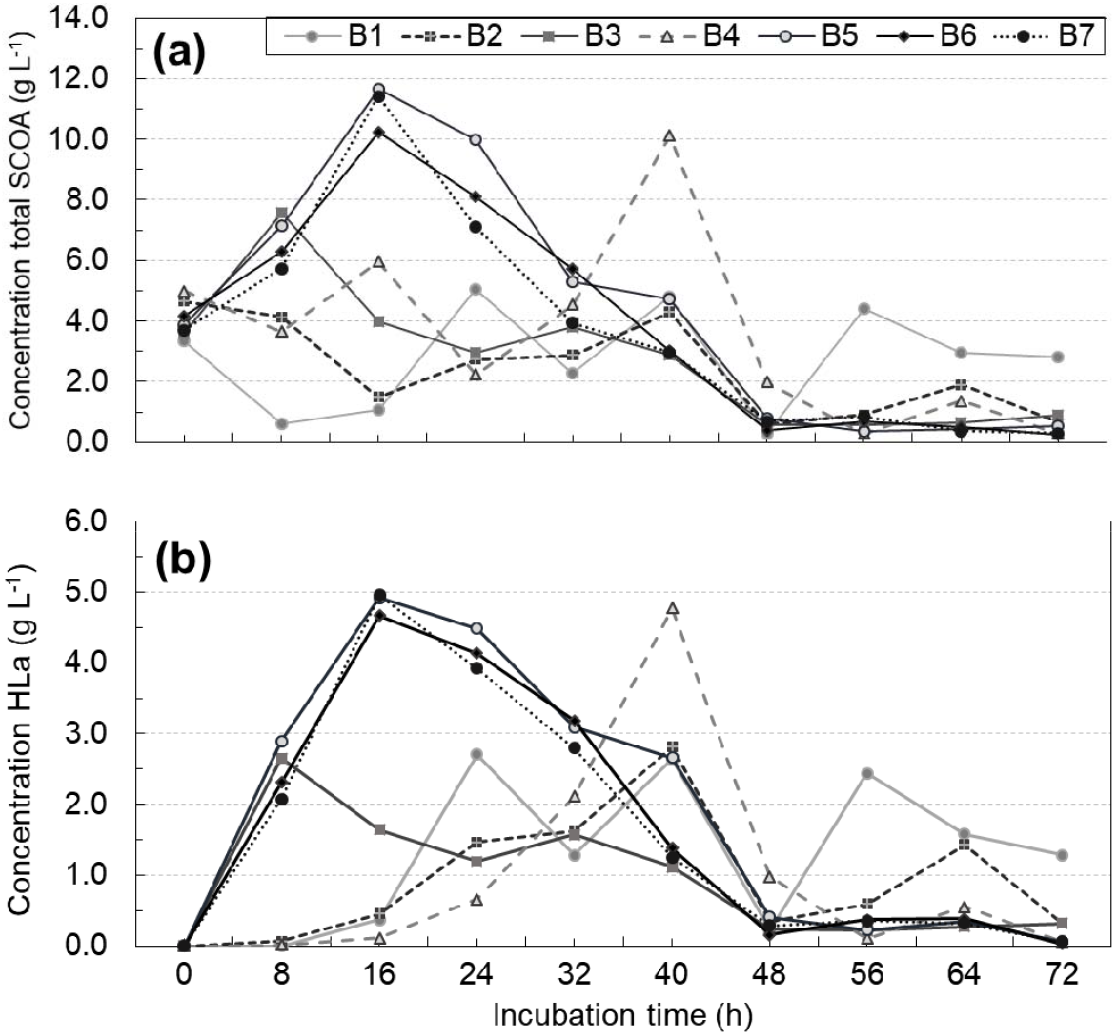
Concentration profiles in batch reactors along 72 h of total operation: (a) total SCOAs and (b) lactic acid (HLa).

As can also be seen in Fig. 2b, reactors B2 and B4, where initial COD of substrate was the highest one (30 g COD L^−1^), also showed initial increasing profiles of lactic acid concentration, although during a longer incubation period, from 0 h to 40 h, attaining maximum of 2.82 and 4.77 g HLa L^−1^ at pH 3.7 and 4.8, respectively. This indicates a less pronounced generation trend of SCOAs at lower pH values, generally below 4.5, which is in accordance with some previous results from technical literature (HAUS *et al*., 2011; LI *et al*., 2014).

Concentration profiles of lactic (HLa), propionic (HPr), isobutyric (HIBu) and butyric (HBu) acids, which were metabolites produced at the most appreciable amounts in batch reactors, are presented in Fig. 3. However, among all SCOAs generated, lactate was the one detected at higher concentrations along all batch tests, while isobutyrate showed the lowest concentrations in majority of incubation times. Besides these compounds, malic (HMa), succinic (HSu), formic (HFo), isovaleric (HIVa) and valeric (HVa) acids were also quantified, although at very small concentrations, and have also contributed to the summation resulting in total SCOAs.

**Fig. 3 –.**
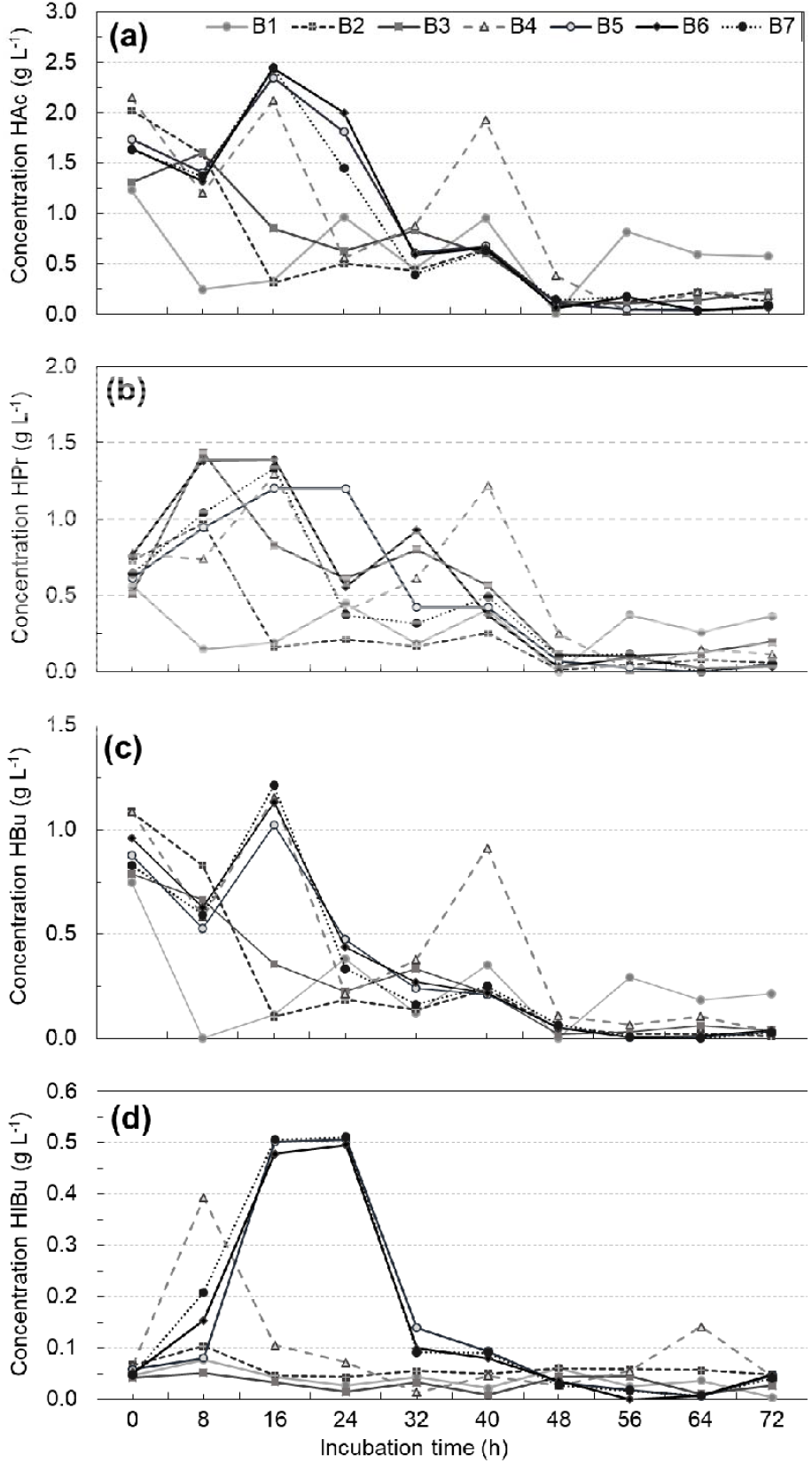
Concentration profiles in batch reactors during 72 h of total operation: (a) acetic acid (HAc), (b) propionic acid (HPr), (c) butyric acid (HBu) and (d) isobutyric acid (HIBu).

In reactors B1 and B3, where initial concentrations of synthetic vinasse were the lowest ones (20 g COD L^−1^), maximum of lactic acid contents were very near each other, 2.71 and 2.66 g HLa L^−1^, occurring at 24 h and 8 h, respectively, see Fig. 2b. In view of this, higher initial pH in reactor B3 seems to have anticipated lactate peak. In accordance to these results, number of SCOAs-producing microorganisms was probably greater at 8 h in reactor B3, which presented an OD 0.145 against an OD 0.126 in reactor B1 at the same incubation time, as shown if Fig. 1c. Additionally, in a general manner, except for reactor B1, all other batch reactors exhibited decreasing lactate concentration profiles right after reaching peaks of concentration at specific incubation times, showing an almost constant behaviour from 48 h to 72 h.

Using food waste from a restaurant as substrate (COD_sol_ ~53 g O_2_ L^−1^), Zhang *et al*. (2005) realized batch tests in order to evaluate influence of pH in hydrolysis and acidogenesis without addition of any extra inoculum (only the ones already existing in food waste were applied to degrade it). In three of four pH levels examined (namely 5, 9 and 11), lactate was obtained at considerable concentrations, often predominant compared to those of formiate, acetate, propionate and butyrate. Similar trends were observed by Wang *et al*. (2001) and Wang (2002), which also assessed anaerobic processing of food wastes using mixed cultures, although both works have focused solely on lactic acid production.

Under pH 5, acidic value that could disfavour the production of SCOAs by *Clostridium* strains (HAUS *et al*., 2011), concentrations of lactic acid (HLa), formic acid (HFo), acetic acid (HAc), propionic acid (HPr) and butyric acid (HBu) attained by Zhang *et al*. (2005) were 14.9 g HLa L^−1^, 1.7 g HFo L^−1^, 1.0 g HAc L^−1^, 4.5 g HPr L^−1^ and 18.5 g HBu L^−1^. In contrast, under pH 11, which could stimulate the generation of SCOAs using *Clostridium* (MONOT *et al*., 1984), Zhang *et al*. (2005) quantified the following concentrations: 20.5 g HLa L^−1^, 8.8 g HFo L^−1^, 13.2 g HAc L^−1^, 1.6 g HPr L^−1^ and 4.8 g HBu L^−1^. Taking these data into consideration, comparing to values observed at 16 h in profiles of Figs. 2a and 3b for reactors B5, B6 and B7 (or at 40 h for reactor B4), for instance, it can be noted there is a certain compatibility between behaviours concerning lactate and acetate, despite they have been generated at considerably higher concentrations by Zhang *et al*. (2005), which applied a strategy of pH control.

Sydney et al (2014) performed anaerobic digestion batch experiments using sugarcane vinasse as carbon source aiming the production of H_2_ and SCOAs by applying two bacteria consortia: one from a sample of fruit bat feces (LPB AH1) and another from a sample of a stabilization lake of a dairy farm (LPB AH2). In preliminary tests, utilization of pure vinasse resulted in low biogas production, so supplementation with sugarcane juice, molasse or sucrose was realized in order to stimulate a greater H_2_ generation. By means of an experimental design to evaluate effects of pH and carbon source content, it was verified that lactate was the major metabolite in most of runs, especially in those carried out with vinasse supplemented with sugarcane juice at concentrations higher than 10 g sugar L^−1^ and using bacteria consortia LPB AH1.

For example, for vinasse mixed to sugarcane juice at 20 g sugar L^−1^ and under pH 5.5, Sydney *et al*. (2014) obtained the following variations in SCOAs contents (generation “+” and consumption “–”): +8.3 g HLa L^−1^, +3.2 g HFo L^−1^, +0.7 g HAc L^−1^, –0,4 g HPr L^−1^ and +2.3 g HBu L^−1^, all of them computed through absolute difference taking non-fermented medium as reference. Comparing these data to those of Fig. 2, one can note there is an agreement regarding lactate has been the metabolite produced at higher concentrations, which was observed in reactor B5, B6 and B7 in period 0-32 h and in reactor B4 in period 0-40 h.

Pursuing generation of H_2_ in a continuous fermentative reactor system using *Enterobacter aerogenes* E. 82005 as microorganism and molasse of unknown origin (13.8 g Suc L^−1^ + 1.5 g Glu L^−1^ + 2.1 g Fru (fructose) L^−1^) as substrate, Tanisho and Ishiwata (1994) obtained lactic acid as the most abundant liquid product, probably by means of a metabolic route similar to the one that seems to have been followed in batch reactors B1-B7 as indicated by profiles of Fig. 2, however not exactly in the same way.

Aiming to evaluate the influence of partial pressure of H_2_ on SCOAs generation, Zhou *et al*. (2017) investigated three innovative methods, namely removal of H_2_ from headspace (T1), aspersion of CO_2_ (T2) and aspersion of H_2_:CO_2_ (80:20), besides the typical method, which consisted in not removing H_2_ as a control test (T4). A glucose solution at 20 g Glu L^−1^ was used as substrate and sludge from a wastewater treatment plant as inoculum for batch reactors. Irrespective of method, lactate was detected as the prevailing metabolite, reaching concentrations of 8.15, 9.85, 6.63 e 7.81 g HLa L^−1^ trough methods T1, T2, T3 e T4, respectively, which is in qualitative accordance with profiles Figs. 2a and 2b. However, formiate, which was not obtained in batch experiments of the present work, were obtained by Zhou *et al*. (2017) in the same way as for Zoetemeyer *et al*. (1982) while they had studied the effect of pH on limited concentration glucose solution (1% Gli) acidogenesis.

#### 3.1.2 Other relevant SCOAs (acetate, propionate, butyrate and isobutyrate) concentrations

Concerning acetic, propionic and butyric acids, batch reactors have been started under average concentrations near to the ones for these compounds in synthetic vinasse for different values of COD_sol_, see Table 2. Particularly for acetic acid, according to Fig. 3a, average initial concentrations were 1.27 g HAc L^−1^ in reactors B1 and B3, 2.09 g HAc L^−1^ in reactor B2 and B4, and 1.67 g HAc L^−1^ in reactors B5, B6 and B7.

For reactor B1, as shown in Fig. 3a, there was consumption of acetic acid during the first 8 h until concentration reaches 0.25 g HAc L^−1^. Differently, in period from 8 h to 24 h, it was observed an increase up to 0.97 g HAc L^−1^. After an alternation of consumption and generation periods, acetate content decreased to zero at 48 h, however it was once gain detected already at 56 h. Finally, a greater stability of lactate concentration was reached between 56 h and 72 h compared to prior incubation time intervals. Also according to Fig. 3a, in opposition to what was observed for reactor B1, reactor B3 presented a production of acetic acid in the first 8 h of up to 1.6 g HAc L^−1^. Subsequently, there was prevalence of consumption periods of acetate until 48 h, when a more defined invariability was initiated until 72 h.

In Fig. 3a it is also observed that acetate content in reactor B2 presented a reduction to 0.31 g HAc L^−1^ from 0 h to 16 h, which was followed by an increase up to 0.64 g HAc L-1 at 40 h. Subsequently, there was consumption of acetic acid between 40 h and 48 h and finally a greater stability from 48 h to 72 h. In reactor B4, however, acetate content decreased to 1.2 g HAc L^−1^ in the first 8 h, while an increase to 2.1 g HAc L^−1^ already occurred at 16 h. Period of 16-48 h was characterized by the intercalation of great consumptions and generations of acetic acid, while period of 48-72 h was of relatively lesser variations.

Still concerning acetic acid, reactors B5, B6 and B7, starting from identical conditions of pH and COD, showed concentration profiles qualitatively similar, see Fig. 3a. In all three reactional media there was consumption of acetic acid in period from 0 h to 8 h, with concentrations attaining 1.40, 1.32 and 1.37 g HAc L^−1^, respectively. In the following incubation times, after peaks have been detected at 16 h, considerable decreases were observed, particularly from 16 h to 48 h. In a general manner, except for reactor B3, all others exhibited reductions in acetate content until 8 h and a more or less well established invariability of it from 48 h to 72 h.

Besides batch anaerobic experiments for SCOAs production, Zhou *et al*. (2017) have also identified some microorganisms existing in fermentative media and found at least twelve genders, with predominance of *Proteus, Enterococcus* and *Clostridium* as the most abundant ones (the two first in methods T1, T2 and T4 and the latter one in method T3). Thus, Zhou *et al*. (2017) have proposed that methods T1, T2 and T4 caused lactic acid generation by means of microorganisms belonging to *Enterobacteriaceae* family, which promoted glucose breakdown reaction into metabolites, resulting in 2 ATP. Additionally, for method T4, Zhou *et al*. (2017) linked the presence of *Clostridium* to the greater acetate content compared to other methods. Thus, it is possible that some microbial genera detected by Zhou *et al*. (2017) also existed in the cattle manure sludge utilized in this work since lactate and acetate were both produced in high appreciably concentrations in batch experiments B1-B7.

Concerning propionic acid, operation of reactors were started from the following average concentrations: 0.54 g HPr L^−1^ for reactors B1 and B3, 0.75 g HPr L^−1^ for reactors B2 and B4, 0.67 g HPr L^−1^ for reactors B5, B6 and B7, as shown in Fig. 3b. Reactors B1 and B3 exhibited a pronounced difference between peaks of propionate concentration, which were 0.45 g HPr L^−1^ at 24 h and 1.43 g HPr L^−1^ at 8 h, respectively. Therefore, reactional medium under higher initial pH (7.5 for reactor B1, against 4.5 for reactor B3) showed the greatest maximum of propionate content, suggesting that it may be advantageous to carry out acidogenesis at higher initial pH (VENTURA e JAHNG, 2013; HAUS *et al*., 2011; LI *et al*., 2014; MONOT *et al*., 1983; MONOT *et al*., 1984). In a similar way, reactors B2 (initial pH 4.5) and B4 (initial pH 7.5) reached maximum of propionate content of 0.97 and 1.29 g HPr L^−1^ at 8 h and 16 h, respectively, without anticipating of greatest concentration peak due to higher initial pH in reactor B4. Additionally, reactors B5, B6 and B7 have attained the greatest propionate concentrations (1.20, 1.39 and 1.34 g HPr L^−1^, respectively).

Isobutyric acid presented similar trends to those for acetic and propionic acids. Thus, reactors under higher initial pH and same initial COD showed greater peaks of isobutyrate concentration. Moreover, as one can see in Fig. 3d, reactors B5, B6 and B7 presented the highest maximum of isobutyric acid content, which were 0.51, 0.50 and 0.51 g HIBu L^−1^, respectively, all occurring at incubation time 24 h.

Regarding butyric acid, its concentration profiles are shown in Fig. 3c. Its average initial concentrations were 0.77 g HBu L^−1^ for reactors B1 and B3; 1.09 g HBu L^−1^ for reactors B2 and B4; 0.89 g HBu L^−1^ for reactors B5, B6 and B7. Maximum butyrate concentration was 0.38 g HBu L^−1^ at 24 h in reactor B1, being lower, however, than the initial content of this compound. In reactor B3, consumption and stability periods were more frequent during batch, excepting the interval 24-32 h, where an increase of 0.11 g HBu L^−1^ was verified, although it was clearly negligible compared to intervals of reductions in butyrate content. In reactors B2 and B4, peaks of butyric acid concentration were 0.24 and 1.16 g HBu L^−1^ at incubation times 40 h and 16 h, respectively. In reactors B5, B6 and B7, maximum of butyric acid concentration were 1.02, 1.13 and 1.21 g HBu L^−1^, respectively, all of them at incubation time 16 h, followed by intervals of consumption and finally of almost invariability.

With the purpose of generating H_2_, Fang *et al*. (2006) carried out fermentation batch experiments of rice slurry as carbon source and sludge from a municipal sewage treatment plant as mixed culture of microorganisms (thermally pre-treated at 100 °C for 30 min). Effects of pH, temperature and carbohydrate (Carb) concentration were assessed. Particularly, evaluation of influence of carbohydrate concentration (2.7-22.1 g Carb L^−1^, pH 4.5 and temperature 37 °C) showed that acetic and butyric acids were the major metabolites, while methanol, propanol, propionic, valeric acid (HVa) and caproic acid (HCa) were also detected, but at relatively lower contents. For example, under 22.1 g Carb L^−1^, acetate and butyrate concentrations were 28.8% and 70.9% out of ~10.6 g total SCOAs + alcohols L^−1^.

An important aspect approached by Fang *et al*. (2006) was the identification of microorganisms, which indicated a greater abundance of *Clostridium* bacteria genus of strictly anaerobic class in sludge. A number of acidogenic species able to produce H_2_ were found, such as *C. acetobutylicum, C. butylicum, C. pasteurianum*, etc., that might as well be present in the cattle manure sludge used in this work, although it cannot be assertively stated based only on profiles of batch reactors B1-B7. Nonetheless, since final reactional medium obtained by Fang *et al*. (2006) contained mainly acetate, butyrate and small amounts of alcohols (besides other minority acids), it is observed disagreement with profiles depicted in Figs. 2 and 3, indicating that distinct routes were followed, which can certainly be associated to different conditions defined for sets of experiments of each work.

Zong *et al*. (2009) executed a process consisting of consecutive dark fermentation and photofermentation in batch mode and using cassava wastewater (18 g hexose L^−1^), food wastes from coffee shop (20 g L^−1^) and sucrose (18 g L^−1^) as carbon sources plus several minerals for composing reactional medium. It is important to note that effluent of dark fermentation was centrifuged and diluted and had its pH adjusted to 7.0 in order to be subsequently processed by photofermentation. A cattle manure compound was applied as inoculum in dark fermentation and isolated cells of *Rhodobacter sphareoides* ZX-5 in photofermentation.

Focus of Zong *et al*. (2009) was the production of H_2_, which was obtained simultaneously to the generation of SCOAs such as acetate, propionate, butyrate and isobutyrate. For instance, fixing initial pH 6.8 (no control), experiment using coffee shop wastes processed by means of dark fermentation provided contents of 2.54 g HAc L^−1^,0.37 g HPr L^−1^, 5.46 g HBu L^−1^, 0.88 g HIBu L^−1^, attaining final pH 4.84. In the same experiment, butanol was obtained at 1.26 g BuOH L^−1^. So, although at lower concentrations, all acidic metabolites detected by Zong *et al*. (2009) were also verified in batch tests of this work, as can be seen in profiles showed in Figs. 2 and 3, while butanol were not produced. This does not exclude definitely the possibility of existence of microorganisms producing butyl alcohol in cattle manure sludge used in reactors B1-B7, however indicates clearly that at least they were not effective to convert substrate in solvent.

By means of batch experiments, Silva *et al*. (2013) evaluated anaerobic acidification of eight different organic materials, namely cheese whey, sugarcane molasse, organic fraction of municipal solid wastes, glycerol, soapy slurry, winery wastewater, olive mill effluent and landfill leachate. Sludge from a treatment plant for domestic and industrial effluents was used as inoculum. Particularly, cheese whey (COD_sol_ 98 g L^−1^), sugarcane molasse (COD_sol_ 896 g L^−1^) and organic fraction of municipal solid wastes (COD_total_ 30 g L^−1^) showed the greatest acidogenic potential, producing 0.33-0.42 g SCOAs (g COD feed)^−1^, with prevalence of acetic, butyric and propionic acids, which were also obtained in batch reactors of the presented work, see Fig. 3, even though they were not the major metabolites in ascending order. However, by varying the proportion substrate/microorganism and alkalinity of medium through addition of CaCO_3_, Silva *et al*. (2013) were able to increase production up to 0.63 g total SCOAs (g COD feed)^−1^, shifting the content predominance to propionic and valeric acids.

Davila-Vazquez et al. (2011) performed batch experiments with cheese whey as substrate (solubilized lactose (Lac) content of 16.5 g Lac L^−1^) and sludge from an UASB reactor for treatment of candy factory wastewater as inoculum. Carbonate- and phosphate-based buffering media were employed in order to reduce pH variations. In tests with carbonate-based buffer carried out in a 3 L fermenter, concentrations of 2.3 g HAc L^−1^, 3.4 g HPr L^−1^, 5.0 g HBu L^−1^ and 0.08 g EtOH (ethanol) L^−1^ were attained through a carbohydrate consumption >99% of available lactose, greater than that in batch reactors of the present work shown in Fig. 1a.

Using a substrate containing lactose (same monosaccharide of cheese whey), Collet *et al*. (2003) observed the presence of lactate in medium resulting from batch fermentation with *Clostridium thermolactitum*. However, opposing to this observation, Davila-Vazquez *et al*. (2011) did not obtained lactate, which probably occurred due to the absence of microorganisms promoting lactic fermentation in mixed culture used as inoculum. Moreover, by comparing results presented here to those by Davila-Vazquez *et al*. (2011), once again one can see quantitative disagreement since ascending order of metabolites concentrations for both works are distinct, even though most of acids produced are common (excepting lactate and isobutyrate).

Additionally, Davila-Vazquez *et al*. (2011) also carried out analyses of microbial communities that have developed in 120 cm^3^ vials containing buffering mineral media. Under the evaluated conditions, results are suggestive of existence of bacteria from *Clostridium* and *Enterobacter* genera, known as efficient H_2_ producers. In this sense, in batch experiments using xylose and cellulose as substrates separately, Lin and Hung (2008) have found microorganisms from *Klebsiella, Pseudomonas, Clostridium* and *Streptococcus* genera in cattle manure sludge. Therefore, despite the similarity of origin of inocula used in Lin and Hung (2008) and in this work (note that specific cattle kind and and it its food diet are also relevant factors), Lin and Hung (2008) have detected other major metabolites, namely H_2_, ethanol, acetic acid, propionic acid and butyric acid.

### 3.2 Maxima of SCOAs productivities

Acidogenesis profiles for batch reactors B1-B7 pointed out that metabolites presenting the highest concentration peaks were lactic, propionic, acetic and isobutyric acids in ascending order. Maxima of productivities of these compounds occurred at specific incubation times and are shown in Table 4. Among all acidic products, lactate exhibited the greatest peak of productivity, observed in reactor B3 at 8 h, which was 332.10 mg HLa L^−1^ h^−1^. However, at the same time and reactor, lactate yield was 189.14 mg HLa (g Glu)^−1^, lower than those calculated for some other batch reactors, indicating that maximum of both performance parameters were not verified simultaneously for lactic acid.

**Table 4 –.** Maxima of productivities and corresponding yields for major metabolites generated in synthetic vinasse acidogenesis for all batch reactors.

| Batch reactor | Incubation time | Lactic acid (HLa) |  | Incubation time | Propionic acid (HPr) |  | Incubation time | Acetic acid (HAc) |  | Incubation time | Isobutyric acid (HIBu) |  |
| --- | --- | --- | --- | --- | --- | --- | --- | --- | --- | --- | --- | --- |
|  |  | *P* | *Y* |  | *P* | *Y* |  | *P* | *Y* |  | *P* | *Y* |
|  | h | mg L<sup>-1</sup> h<sup>-1</sup> | mg (g Suc<sup>-1</sup>) | h | mg L<sup>-1</sup> h<sup>-1</sup> | mg (g Suc<sup>-1</sup>) | h | mg L<sup>-1</sup> h<sup>-1</sup> | mg (g Suc<sup>-1</sup>) | h | mg L<sup>-1</sup> h<sup>-1</sup> | mg (g Suc<sup>-1</sup>) |
| B1 | 24 | 112.79 | 205.93 | *– | – | – | – | – | – | 8 | 3.89 | 2.36 |
| B2 | 40 | 70.40 | 146.81 | 8 | 30.30 | 12.64 | – | – | – | 8 | 4.63 | 1.93 |
| B3 | 8 | 332.10 | 189.14 | 8 | 115.28 | 65.65 | 8 | 36.39 | 20.72 | 8 | 1.20 | 0.69 |
| B4 | 40 | 119.32 | 233.09 | 16 | 32.48 | 25.38 | – | – | – | 8 | 40.65 | 15.88 |
| B5 | 16 | 307.52 | 303.40 | 8 | 40.90 | 20.18 | 16 | 38.08 | 37.57 | 16 | 27.72 | 27.35 |
| B6 | 16 | 291.54 | 299.77 | 8 | 77.58 | 39.88 | 16 | 49.74 | 51.15 | 16 | 26.70 | 27.45 |
| B7 | 16 | 309.88 | 306.30 | 8 | 49.37 | 24.40 | 16 | 50.84 | 50.25 | 16 | 28.65 | 28.32 |

Labels: \*symbol “–” indicates there was consumption of metabolite during operation of batch reactor regarding initial value of concentration (incubation time 0 h).

Reactors B5, B6 and B7, all prepared under identical initial pH and COD, presented maximum of lactic acid productivity at 16 h, although value obtained for this parameter was slightly higher for reactor B7 (309.88 mg HLa L^−1^ h^−1^). Since incubation time of 0 h (starting point for experiments) was taken as a reference to compute *P* and *Y*, see Eqs. (1) and (2), a similar trend to that for productivities applies for yields in reactors B5, B6 and B7 as well, with a very small advantage for reactor B7, which presented a little bit higher maximum of lactate yield than others (306.30 mg HLa (g Glu)^−1^).

In all batch reactors (B1-B7), peaks of productivity and corresponding yields of lactic acid were expressively greater than those of other metabolites. Invariably, as shown in Table 4, the highest maximum of productivity for propionate, acetate and isobutyrate were observed at incubation times of 8 h and 16 h. For propionic acid, maximum of productivity was 115.28 mg HPr L^−1^ h^−1^ at 8 h, with a simultaneous yield of 65.65 mg HPr (g Glu)^−1^. For acetic and isobutyric acids, maximum of productivity and yield were limited up to about 50 g L^−1^ h^−1^ and 50 mg (g Glu)^−1^.

By means of anaerobic digestion batch experiments also employing cattle manure sludge as inoculum, Moraes *et al*. (2019) obtained considerably high concentrations of lactic acid mainly when using a mixture of real vinasse and a synthetic culture medium at 1:3 v/v. Actually, lactate was the major metabolite in these specific tests, regardless of pre-treatment method applied to sludge. However, tests performed under higher contents of real vinasse in mixture, particularly at 2:1 v/v, showed profiles indicating there was consumption of lactate in certain incubation periods. Additionally, in the same way as Lappa *et al*. (2015b), Moraes *et al*. (2019) linked the increase in butyrate concentration to the reduction of lactate concentration.

Despite of Moraes *et al*. (2019) have not presented lactic acid productivities and yields probably because lactate was not taken as a product of interest of acidogenesis, concentrations and incubation times were given, namely a maximum of 8.8 g HLa L^−1^ at 144 h (6 days) from an initial carbohydrate content of 20 g Carb L^−1^. In view of this, it can be computed via Eq. (1) that peak of productivity was 61.1 g HLa L^−1^ h^−1^, resulting in a yield 440.0 mg HLa (g Carb)^−1^ by Eq. (2). Both values of parameters were verified at 1:3 v/v with sludge being pre-treated by acidic + thermal method. By comparing these results to those shown in Table 4, one can presume that lower incubation time associated to maximum of productivity of present work (8 h, 332.10 g HLa L^−1^ h^−1^) is the cause of difference.

Using vinasse as substrate, Djukić-Vuković *et al*. (2015) conducted lactic fermentation experiments in presence of *Lactobacillus rhamnosus* ATCC 7469 strain and have attained a maximum of 50.18 g HLa L^−1^ for lactate concentration, resulting in a corresponding productivity of 1480 mg HLa L^−1^ h^−1^ and a yield of 900 mg HLa (g Carb)^−1^. Comparing them to those listed in Table 4, the greatest values of performance parameters obtained by Djukić-Vuković *et al*. (2015) are certainly due to use of a *Lactobacillus* species, highly specialized in producing lactate, as fermentative microorganism instead of a mixed culture containing a wide microbial diversity, which was the case of present work.

Using vinasse as nutrient source and cells of *Lactobacillus rhamnosus* ATCC 7469 fixed over zeolite particles as inoculum, Djukić-Vuković *et al*. (2013) attained a maximum of 42.19 g HLa L-1 for lactate content, with a correspondent productivity of 1690 g HLa L^−1^ h^−1^ and yield of 960 g HLa L^−1^. Employing the same microorganism, but this time deposited on the surface of Mg-modified zeolite particles, Djukić-Vuković *et al*. (2016) reached maximum of 47.60 g HLa L^−1^ (lactate content), 1410 mg HLa L^−1^ h^−1^ (lactate productivity) and 860 g HLa L^−1^ (lactate yield). Taking into consideration both these previous studies, lactate concentration and performance parameters were higher than those detected in the present work because carbohydrate content in substrate was notably greater, 50 g Carb L^−1^ (DJUKIĆ-VUKOVIĆ *et al*., 2013; DJUKIĆ-VUKOVIĆ *et al*., 2016), compared to a maximum concentration of 18 g Glu L^−1^ adopted for synthetic vinasse in batch experiments B1-B7. Moreover, *Lactobacillus rhamnosus* ATCC 7469 is surely more suitable for lactic acid generation than cattle manure sludge utilized here.

### 3.3 Possible biogas generation and process mass balance

Anaerobic digestion performed by mixed cultures of bacteria is certainly one of the most common ways of producing SCOAs from a great number of substrates, with emphasis on residual materials. In many cases, however, the goal of acidogenic anaerobic digestion is not exactly the generation of SCOAs, which presents a wide range of applications, but the generation of biogas since chemical reactions resulting in butyric and acetic acids from a carbon source also results in H_2_, a compound with interesting properties for use as gaseous fuel. These reactions are shown in Eqs. (6) and (7) for glucose, a directly fermentable sugar, as substrate.

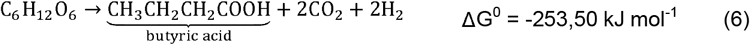

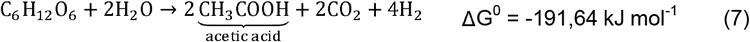

In chemical reactions producing lactic and propionic acids (both metabolites that can also be relevant in acidogenesis depending on conditions it is performed) from glucose, see Eqs. (8) and (9), one can note there is no formation of CO_2_ and H_2_, which differs clearly from what is observed in Eqs. (6) and (7). In contrast, as show in Eq. (9), actually there is consumption instead of generation of H_2_ in reaction providing propionic acid.

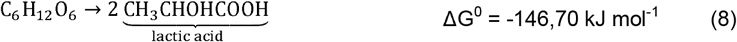

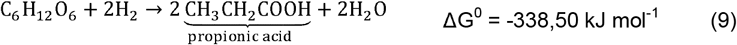

So, even though productive purposes are distinct, it is appropriate to realize comparisons of results of present work, obtained through experiments aiming to generate SCOAs, to results from previous works that focused on biogas production, mainly H_2_. In view of this, results from technical literature and also results of some of batch experiments B1-B7 are reunited in Table 5. Based on them, the most notable observations one can do are the following: absence of lactic acid in majority of fermentative media and prevalence of acetic and butyric acids (the two most common in acidogenesis) in a wide number of references. Additionally, it is important to highlight that ethanol, already existing in original synthetic vinasse at different COD_sol_, did not show significant content variations in tests with cattle manure sludge of the present work.

**Table 5 –.** Concentrations of major SCOAs generated by means of acidogenic fermentation in batch reactors with a diversity of substrates and mixed cultures from different origins.

| Substrate | Mixed culture | pH | Concentration (g SCOA L<sup>-1</sup>) |  |  |  | Reference |
| --- | --- | --- | --- | --- | --- | --- | --- |
|  |  |  | HBu | HAc | HLa | HPr |  |
| Food waste (53.18 g COD<sub>sol</sub> L<sup>-1</sup>) | Microorganisms existing in original substrate | 5.5 (controlled) | 18.5 | 1.0 | 14.9 | 4.5 | Zhang *et al.* (2005) |
| Rice slurry (22.1 g Carb L<sup>-1</sup>)* | Sludge from a municipal sewage treatment plant | 4.5 (controlled) | 7.5 | 3.0 | NA* | NA | Fang *et al.* (2006) |
| Cassava wastewater (18 g Carb L<sup>-1</sup>) | Cattle dung compost (dark fermentation) | 6.8 (initial)<br>4.8 (final) | 4.3 | 2.2 | NA | 0.4 | Zong *et al.* (2009) |
| Sucrose (18 g Carb L<sup>-1</sup>) | Cattle dung compost (dark fermentation) | 6.8 (initial)<br>4.8 (final) | 5.1 | 2.1 | NA | 0.2 | Zong *et al.* (2009) |
| Cheese whey (with a carbonate-based buffering medium) (16.5 g Lac L<sup>-1</sup>) | Sludge from a UASB reactor for treatment of candy factory wastewater | 7.5 (initial)<br>5.4 (final) | 5.0 | 2.3 | NA | 3.4 | Davila-Vazquez *et al.* (2011) |
| Cheese whey (with a phosphate-based buffering medium) (16.5 g Lac L<sup>-1</sup>) | Sludge from a UASB reactor for treatment of candy factory wastewater | 7.5 (initial)<br>4.9 (final) | 1.8 | 1.1 | NA | 2.1 | Davila-Vazquez *et al.* (2011) |
| Sugarcane vinasse (29.6 g COD L<sup>-1</sup>) supplemented with juice (20 g Glu L<sup>-1</sup>) | Sample of fruit bat feces | 5.5 (final) | +2.3 | +0.8 | +8.3 | -0.4 | Sydney *et al.* (2014)** |
| Sugarcane vinasse (29.6 g COD L<sup>-1</sup>) supplemented with molasse (20 g Glu L<sup>-1</sup>) | Sample from a (stabilization) lake of a dairy farm | 5.5 (final) | +3.7 | -0.6 | 0.0 | -0.7 | Sydney *et al.* (2014)** |
| Cheese whey (98 g COD<sub>sol</sub> L<sup>-1</sup>) | Sludge from a treatment plant for domestic and industrial effluents | 6.35 (initial) | ~0.87 | ~1.67 | NA | ~0.24 | Silva *et al.* (2013)*** |
| Sugarcane molasse (896 g COD<sub>sol</sub> L<sup>-1</sup>) | Sludge from a treatment plant for domestic and industrial effluents | 5.83 (initial) | ~0.41 | ~1.60 | NA | ~0.27 | Silva *et al.* (2013)*** |
| Glucose solution (20 g Glu L<sup>-1</sup>) (T2, with CO<sub>2</sub> aspersión) | Sludge from a wastewater treatment plant | ~6.60 (initial)<br>~4.15 (final) | NA | ~0.37 | 9.85 | ~0.10 | Zhou *et al.* (2017)*** |
| Synthetic vinasse (20 g COD L<sup>-1</sup>) (reactor B1) | Cattle manure sludge | 4.5 (initial)<br>4.2 (final) | 0.38 | 0.97 | 2.71 (24 h) | 0.45 | Present work<sup>§</sup> |
| Synthetic vinasse (25 g COD L<sup>-1</sup>) (reactor B7) | Cattle manure sludge | 6.0 (initial)<br>4.8 (final) | 1.21 | 2.45 | 4.96 (16 h) | 1.34 | Present work<sup>§</sup> |
| Synthetic vinasse (30 g COD L<sup>-1</sup>) (reactor B4) | Cattle manure sludge | 7.5 (initial)<br>4.8 (final) | 0.91 | 1.93 | 4.77 (40 h) | 1.22 | Present work<sup>§</sup> |

Labels: NA\*: not available or not detected; \*Higher concentration of substrate adopted by Fang *et al.* (2006); \*\*Presented generation (+) and consumption (-) taking the blank as reference, however did not give concentrations of SCOAs in non-fermented media (blank); \*\*\*Concentrations preceded by the symbol “~” were extracted from graphics, so they are approximations; §Parameters verified at the incubation time of maximum concentration of lactate.

**Table 6 –.** Mass balance in terms of COD at 72 h for all batch reactors.

| COD<br>(10<sup>-3</sup> g L<sup>-1</sup>) | Batch reactors |  |  |  |  |  |  |
| --- | --- | --- | --- | --- | --- | --- | --- |
|  | B1 | B2 | B3 | B4 | B5 | B6 | B7 |
| HMa | 54.72 | 47.17 | 53.75 | 48.42 | 63.62 | 61.42 | 60.49 |
| HSu | 32.97 | 22.82 | 31.15 | 28.19 | 16.08 | 14.69 | 17.88 |
| HLa | 1376.16 | 345.37 | 344.14 | 57.15 | 63.86 | 38.22 | 68.50 |
| HFo | 4.50 | 6.96 | 3.85 | 7.31 | 3.47 | 3.48 | 3.50 |
| HAc | 612.11 | 134.68 | 240.43 | 206.44 | 81.42 | 69.71 | 93.84 |
| HPr | 550.64 | 89.55 | 297.79 | 173.42 | 74.61 | 51.18 | 82.14 |
| HIBu | 6.29 | 88.17 | 48.30 | 81.68 | 85.61 | 86.74 | 73.76 |
| HBu | 386.49 | 20.29 | 70.77 | 62.85 | 56.41 | 72.59 | 52.85 |
| HIVa | 60.42 | 59.13 | 57.05 | 61.17 | 43.74 | 44.14 | 44.92 |
| MeOH | 84.44 | 117.14 | 79.50 | 113.48 | 115.34 | 122.39 | 116.97 |
| EtOH | 878.15 | 737.90 | 321.42 | 331.19 | 115.66 | 108.54 | 129.96 |
| Suc | 5145 | 5675 | 8297 | 9570 | 9777 | 10879 | 11418 |
| Bio | 590 | 1510 | 720 | 742 | 930 | 524 | 585 |
| $COD_{t,T}$ | 9913 | 9.188 | 10724 | 11647 | 11632 | 12191 | 12877 |
| $COD_{exp}$ | 16775 | 24025 | 18025 | 23025 | 21050 | 20550 | 21550 |
| $COD_{exp} - COD_{t,T}$ | 6862 | 14837 | 7301 | 11378 | 9418 | 8359 | 8673 |
| Agreement<br>(Ag, %) | 59 | 38 | 59 | 51 | 55 | 59 | 60 |

It should be emphasized that synthesis reactions of SCOAs, Eqs. (6)–(9), are exergonic since variations of free Gibbs energy for them, estimated by means of e-Quilibrator tool (FLAMHOLZ *et al*., 2011), are all negative. So, these are spontaneous processes, although their degrees of spontaneity are different from each other. Moreover, trends showed in Figs. 2 and 3 sometimes are not in accordance to some cause and effect relationships found in technical literature for process monitoring parameters and concentrations of SCOAs. However, this would be expected because several factors interfering in experimental results were not measured or controlled. In this sense, besides direct comparisons to other works are difficult due to different operating conditions, identification of microorganisms existing in cattle manure sludge would allow a clearer understanding of causes, which however was not carried out in present work. Despite of it, trends verified are useful in some ways, even as preliminary results, mainly to indicate directions favouring production of interest metabolites, although these directions are not definitive.

In profiles showed in Figs. 2 and 3, intervals near to the end of batch experiments are remarkable, particularly after 48 h, when concentrations of SCOAs in most of reactors were notably low, showing periods of consumption of these metabolites. A reasonable explanation for this behaviour seems to be the generation of H_2_, carbon dioxide (CO_2_) and CH_4_, the latter certainly at lower amount. All of them are gaseous products probably generated by microbe existing in cattle manure sludge, but that have not been identified or quantified. Therefore, acidic + thermal pre-treatment may not have been effective to eliminate completely the methanogenic archaeas and these microorganisms may have been developed, even in an incipient way, and performed intensively in periods where contents of acids were decreased, since this is an indication of methanogenesis occurrence. In contrast, for periods where acids contents were increased, acting of SCOAs-producing bacteria, which also generate H_2_ and CO_2_, see Eqs. (8) and (9), was surely more pronounced. In fact, in batch experiments of this work, gas production was verified empirically.

Similarly to results presented here, for anaerobic digestion processes of 1G and 2G (first and second generation) vinasses with sludge from a UASB reactor degrading 1G vinasse, Silverio *et al*. (2019) verified the alternation of periods of generation and consumption of acids throughout feed batch experiments. For periods of accumulation of acids, acetogenic bacteria and hidrogenotrophic and acetoclastic archaeas surely presented small activity (SILVERIO *et al*., 2019). On the other hand, for periods in which there were reductions of acids concentrations, it is reasonable that the opposite justification is valid (MOSEY, 1982; SILVERIO *et al*., 2019). These arguments provide some support to the hypothesis that there was CH_4_ generation in experiments with cattle manure sludge of the present work, although a more categorical evidence is lacking, especially because of acidic + thermal pre-treatment that was performed to inoculum.

Finally, mass balances at 72 h (final incubation time) for all batch reactors are presented in Table 5 and reveals poor agreement between total theoretical and experimental CODs (35-60%), suggesting there were indeed other products in reactional media which were not accounted for. Therefore, it is probable that production of unidentified liquid and dissolved gaseous metabolites has occurred in experiments conducted in this work for assessment of initial pH and COD of synthetic vinasse (PEIXOTO *et al*., 2012; AMORIM *et al*., 2014; MARTINS AND AMORIM, 2016). This could be taken as a consequence of using cattle manure sludge since it certainly contains a number of different microbes capable of generating several compounds undetected as fermentative products.

## 4. CONCLUDING REMARKS

The greatest peaks of total SCOAs concentration were observed at intermediate levels adopted for initial pH and COD, i.e. 6.0 and 25 g COD L^−1^. Maximum of total SCOAs concentration for initial pH and COD of 7.5 and 30 g COD L^−1^ (highest levels of both parameters) were also considerable, although lower than those verified under initial pH 6.0 and 25 g COD L^−1^ and occurring at incubation times further ahead. On the other hand, SCOAs generation seems to have been inhibited in experiments under initial pH 4.5, regardless of COD level (20 or 30 g COD L^−1^), since maximum of metabolites contents in these cases were notably lower compared to those at other conditions.

Based on results of batch tests B1-B7, it was deduced that pH values nearer to neutrality, particularly at the two highest levels (6.0 and 7.5), were determining of occurrence of peaks of SCOAs concentrations. In this sense, even at the highest COD level (30 g COD L^−1^), which would tend to stimulate a greater production of metabolites due to higher content of carbon source available in fermentative medium, maximum of SCOAs concentrations verified under initial pH 4.5 were lower than those observed under initial pH 6.0 and 7.5 combined with the lowest levels of COD, i.e. 20 and 25 g COD L^−1^. Therefore, even at the lowest COD, higher initial pH values were decisive for obtaining greater maximum of total SCOAs concentration.

In most of incubation times, lactate contents were prevailing, suggesting that fermentative process followed an essentially lactic route. Consequently, concentration profiles of total SCOAs showed very similar behaviours to lactate concentration profiles. Aiming to explain this observation, the hypothesis of existence of a large number of microorganisms promoting lactic fermentation in cattle manure sludge was raised. However, it was not possible to accept or reject it definitely based only on obtained results. Moreover, considering that the prepared synthetic vinasse already contained pre-defined amounts of butyrate, acetate and propionate, production of these metabolites was taken as significant only at specific incubation times since their concentrations commonly fluctuated around the initial concentrations during the process, often being reduced to levels below those of the original substrate.

## ACKNOWLEDGMENTS

Authors are grateful to CNPq (Brazilian National Council for Scientific and Technological Development – Finance Code 142340/2016-2) for scholarship and financial support for research and to FAPESP (The São Paulo Research Foundation), Program for Research on Bioenergy (BIOEN) – Regular Program Grants, grant number 2012/09785-8.

